# A Unified Probabilistic Modeling Framework for Eukaryotic Transcription Based on Nascent RNA Sequencing Data

**DOI:** 10.1101/2021.01.12.426408

**Authors:** Adam Siepel

**Affiliations:** Simons Center for Quantitative Biology, Cold Spring Harbor Laboratory, Cold Spring Harbor, NY, USA

## Abstract

Nascent RNA sequencing protocols, such as PRO-seq and NET-seq, are now widely used in the study of eukaryotic transcription, and these experimental techniques have given rise to a variety of statistical and machine-learning methods for data analysis. These computational methods, however, are generally designed to address specialized signal-processing or prediction tasks, rather than directly describing the dynamics of RNA polymerases as they move along the DNA template. Here, I introduce a general probabilistic model that describes the kinetics of transcription initiation, elongation, pause release, and termination, as well as the generation of sequencing read counts. I show that this generative model enables estimation of separate pause-release rates, termination rates, and the initiation/elongation rate ratio up to a proportionality constant. Furthermore, if applied to time-course data in a nonequilibrium setting, the model can be used to estimate elongation rates. This model leads naturally to likelihood ratio tests for differences between genes, conditions, or species in various rates of interest. If read counts are assumed to be Poisson-distributed, convenient, closed-form solutions are available for both parameter estimates and likelihood-ratio-test statistics. Straightforward extensions of the model accommodate variability in the pause site and steric hindrance of initiation by paused polymerases. Additional extensions address Bayesian inference under the Poisson model and a generalized linear model that can be used to discover genomic features associated with rates of elongation. Finally, I address technicalities concerning estimation of library size, normalization and sequencing replicates. Altogether, this modeling framework enables a unified treatment of many common tasks in the analysis of nascent RNA sequencing data.

## Introduction

In recent years, several experimental protocols have emerged for measuring newly produced RNAs on a genome-wide scale [1–5]. These *nascent RNA sequencing* methods can be thought of as adaptations of traditional nuclear run-on assays that exploit modern massively-parallel sequencing technologies to simultaneously map the positions of engaged RNA polymerases across an entire genome. They have proven useful in a wide variety of applications, including measurement of transcription levels independent of RNA decay [1], measurement of rates of elongation [6], mapping of transcription start sites [7], identification of active enhancers [8,9], and estimation of relative RNA half-lives [10]. These measurements have led, in turn, to a number of new insights into the biology of transcription, such as the widespread incidence of promoter-proximal pausing and divergent transcription [1, 2], the remarkably similar architecture of transcription initiation at promoters and enhancers [7], pause-release as a crucial rate-limiting step in transcription [11], and distinct waves of transcriptional responses to the induction of transcription [9].

A variety of computational and statistical methods have been used to analyze these data, ranging from relatively simple count- and ratio-based tests to hidden Markov models and discriminative machine learning approaches (e.g., [6, 8, 9, 12, 13]). Some of these methods are sophisticated, powerful, and widely used. For the most part, however, they have focused fairly narrowly on specific prediction tasks (such as elongation rate estimation or TSS prediction), rather than on more generally modeling the biophysical processes underlying transcription. As a result, with a few partial exceptions [14, 15], the existing computational methods for nascent RNA sequence data, or closely related data types, generally do not permit direct estimation of biophysical parameters of interest, such as rates of initiation or termination. In addition, owing to their heuristic description of the underlying processes, they can be difficult to adapt for nuances in the process, such as promoter-proximal pausing or variation in elongation rate along the gene body.

In this article, I introduce a unified framework that combines a simple kinetic model of transcription with a generative model for nascent RNA sequence data, and permits direct inference of kinetic parameters from sequence data. Separate layers in the model account for stochasticity in the process and noise in the sequencing data, and extension allows for uncertainty in the location of the pause site. I show that this model can be applied to a number of problems of interest, including estimation of initiation and elongation rates. It also leads directly to powerful statistical tests for differences in rates of interest, which naturally consider both the information in the read count data and the underlying physical process. In its simplest form, the modeling framework ignores collisions between RNA polymerase molecules, but I show that it can be extended to allow for certain classes of collisions. In addition, the model has natural extensions to Bayesian inference and to a generalized linear model that accommodates covariates of elongation rate. Finally, I show how to address technical issues concerning estimation of library size, normalization, and sequencing replicates within this modeling framework. I conclude with a discussion of some current limitations and possible extensions of the current framework.

### General Kinetic Model for Transcriptional Initiation, Elongation, Pause-Release, and Termination

I begin with a general kinetic model for the process by which RNA polymerase (RNAP) enzymes initiate transcription at the transcription start site (TSS) of a transcription unit (TU) and move along the DNA template, synthesizing an elongating nascent RNA molecule. This model is defined by a series of states corresponding to each nucleotide position of RNAP as it moves along the template (Figure 1). In particular, state *i* represents the positioning of the active site of RNAP at nucleotide *i* of the TU and corresponds to a nascent RNA of length *i* − 1. I assume that each RNAP eventually traverses the entire DNA template, borrowing from evidence suggesting that premature termination occurs at relatively low rates across most transcription units [16] (but see Discussion).

**Figure 1:**
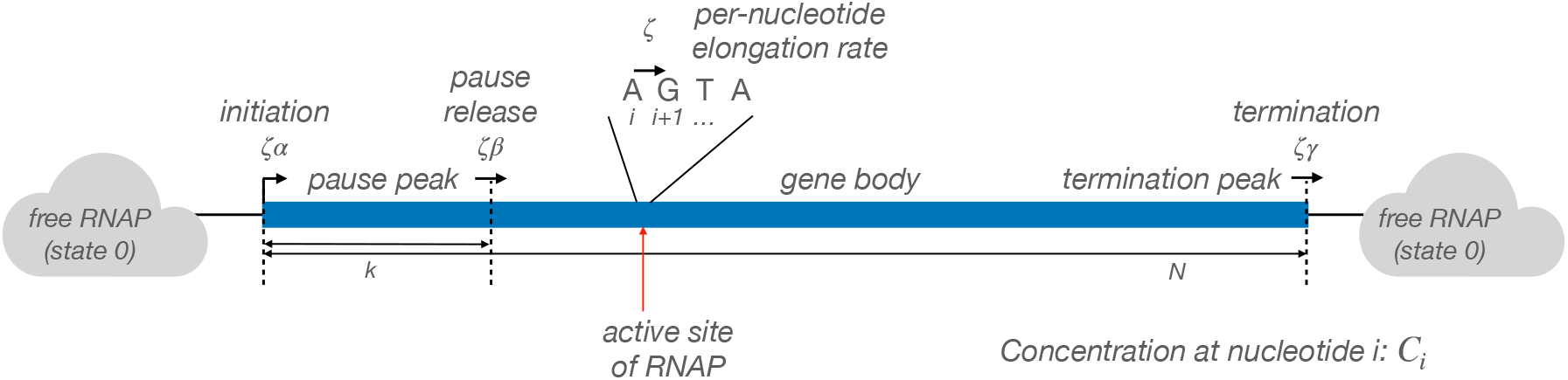
General kinetic model for transcription initiation, elongation, pause release, and termination along a transcription unit.

I will distinguish between two segments of a TU of length *N* : (1) the first *k* nucleotides, known as the *pause peak*, where RNAP tends to accumulate owing to promoter-proximal pausing (typically *k* ≈ 50) [1]; and (2) the subsequent *N − k* nucleotides, where RNAP tends to be relatively unimpeded, which is typically referred to as the *gene body*. I will also allow for a *termination peak* near the end of the TU, owing to delays from termination of transcription (see Figure 2). The model has *N* + 1 states, corresponding to the *N* nucleotide positions of the TU plus an additional state (labeled 0) that abstractly represents *free RNAP*, that is, RNAP that is not currently engaged in transcription and is available for a new initiation.

**Figure 2:**
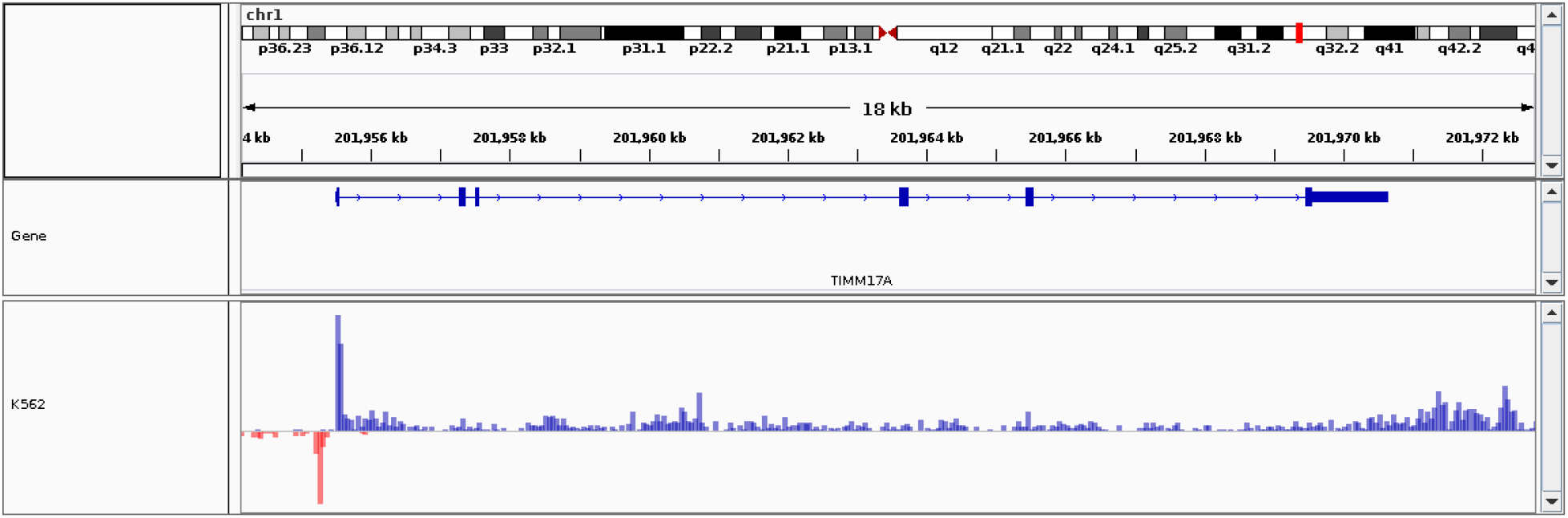
Example of real nascent RNA sequencing data in the region of the human *TIMM17A* gene. The data were obtained by applying PRO-seq to K562 cells, as described in ref. [9]. The pause peak and termination peak are clearly visible. Notice, however, that the read counts are quite noisy along the gene body. In addition, transcription in this case seems to have run past the annotated transcription termination site, a phenomonen that occurs frequently. As a result, the methods described in this paper would likely require some adjustment of annotated gene coordinates based on the raw data itself (e.g., see [13]).

I assume, as with most existing protocols, that the data summarize a relatively large population of cells, and denote by *C_i_* the concentration of cells in the sampled population having RNAP positioned at nucleotide *i*. As will be seen below, the sequencing reads from a PRO-seq experiment that map to position *i* at their 3′ ends should have a depth roughly proportional to *C_i_*. The situation is similar for other protocols.

The kinetic model is defined by four rates: an initiation rate *α*, a pause-release rate *β*, a termination rate *γ*, and a constant per-nucleotide elongation rate *ζ*. For mathematical convenience, I assume that the initiation, pause-release, and termination steps are coupled with single-nucleotide elongation steps and occur at rates *ζα*, *ζβ*, and *ζγ*, respectively. As a result, as long as *ζ* is the same across nucleotides, it can be considered a scaling factor that applies equally to all steps in the process. (Later I will consider a generalization that allows this rate to vary across nucleotides.) I will also assume, for now, that the RNAPs are relatively sparse along each DNA template, and ignore the effects of collisions between them, although this simplification, too, will be revisited.

With these assumptions, the concentrations *C_i_*, for *i* ∈ {0, …, *N*}, are governed by the following system of *N* + 1 differential equations,

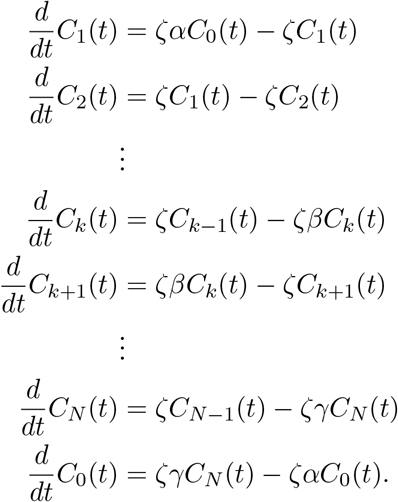

I will consider both the time evolution of the *C_i_* values and their values at steady-state.

### Continuous-time Markov Model

If instead of thinking in terms of a population of cells, one considers the probability distribution over states (nucleotide-specific positions) for a given RNAP, the same model can be represented using the convenient and flexible formalism of continuous-time Markov models. The continuous-time Markov version of the model consists of *N* + 1 states and transition rates exactly analogous to the ones above (Figure 3). In this case, however, each state *i* corresponds to a binary random variable *Z_i_*, indicating whether a given RNAP is (*Z_i_* = 1) or is not (*Z_i_* = 0) at a particular position at a particular time *t*. The assumption that collisions are rare allows the stochastic process followed by each RNAP to be considered independent of all of the others.

**Figure 3:**
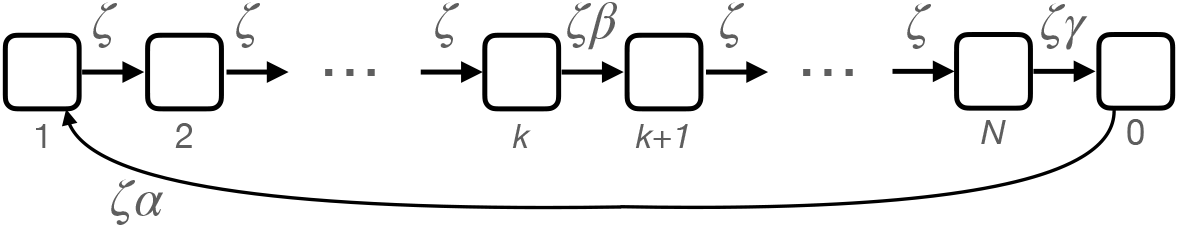
State-transition diagram for continuous-time Markov model for a single RNAP molecule moving along the DNA template.

It is easy to see that this model is identical to the previous one up to a normalization term. Given a sufficiently large collection of independent RNAP molecules, the overall concentration *C_i_*(*t*) is expected to be proportional to the probability of occupancy *P*(*Z_i_*|*t, α, β, γ, ζ*) for any individual RNAP. Furthermore, for *P*(*Z_i_*|*t, α, β, γ, ζ*) to remain a proper probability distribution, the constant of proportionality must be given by 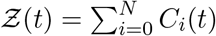.

This Markov chain is defined by an (*N* + 1) × (*N* + 1) infinitesimal generator matrix, **Q** = *ζ***R**, such that,

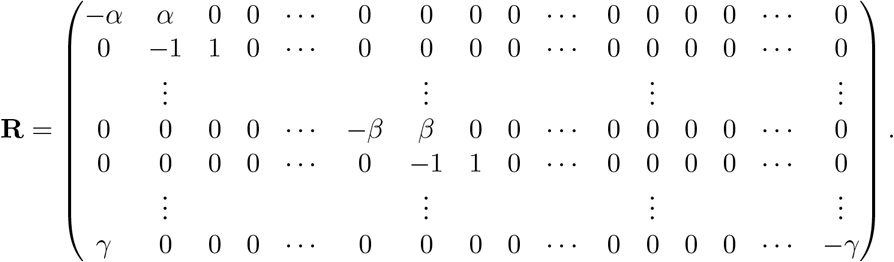

The element at row *i* and column *j* of **Q** indicates the instantaneous rate at which transitions occur from state *i* to state *j*. By convention, the values along the diagonal are set such that the rows sum to zero. Because the states must be visited in a sequence, this matrix has a particularly simple form: it has nonzero terms only on the main diagonal, the diagonal immediately above it, and the single element at bottom left (which allows RNAPs to “wrap around” from the last state to the first one).

The probability of transitioning from any state *i* to any other state *j* over a period of time *t* ≥ 0 can be computed in the usual way as,

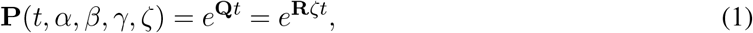

where the element at row *i* and column *j* of the matrix **P**() indicates the conditional probability of occupancy of state *j* at time *t* given occupancy of state *i* at time 0, and the matrix exponential is given by the Taylor expansion 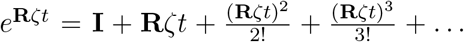 [17]. If transcription is initiated at time 0 (meaning that the initial state is state 0), then the distribution over states at a later time *t* is simply given by row *i* = 0 of **P**(). In particular, the element at the *j*th column of row *i* = 0 indicates the quantity I have denoted *P*(*Z_j_|t, α, β, γ, ζ*).

Notice that the elongation rate *ζ* and the time *t* are nonidentifiable under these assumptions; one of these quantities must be known to estimate the other.

As discussed further below, it will sometimes be desirable to allow for uncertainty in *k*, according to some prior distribution *P*(*k*). In this case, the transition matrix can simply be defined as a mixture

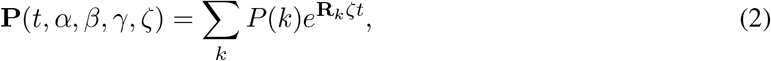

where each **R**_*k*_ reflects a different choice of *k*.

### Generative Model for Sequence Data

In the analysis of nascent RNA sequencing data, we have the additional complication that the positions of RNAP molecules are not directly observed, but instead are indirectly indicated by aligned sequence reads. The read counts reflect a sampling process that probes only a relatively small subset of cells in the starting material. Moreover, this process can be biased by factors such as base composition or RNA secondary structure that can lead to subtle differences in the efficiency of various steps in library preparation, including DNA fragmentation, purification, and PCR amplification.

I will address this problem by adding a second layer to the model that describes the probabilistic process by which sequencing read counts are generated conditional on an underlying “true” density of RNAP at each nucleotide. In this way, I can obtain a full generative model for the observed sequence data that is defined by the parameters of both the kinetic model described above and these conditional distributions for sequencing reads, enabling inference of both sets of parameters from data.

Let *μ_i_* be the expected read depth at position *i*. I will assume that *μ_i_* is proportional to *P*(*Z_i_|t, α, β, γ, ζ*) (as computed from the continuous-time Markov chain), with a proportionality constant that depend on the sequencing depth, that is, with *μ_i_* = *λP*(*Z_i_|t, α, β, γ, ζ*), where *λ* is a measure of sequencing depth. In addition, I will abstractly assume a general generating distribution, *ψ*, for read counts given *μ_i_* and any other relevant parameters, which will be cumulatively denoted *θ*. Let the data be denoted **X** = (*X*_1_, …, *X_N_*), where *X_i_* representes the number of sequencing reads having their 3′ end aligned to position *i*. Assuming independence of sequencing reads, the likelihood for the data in terms of the time *t* and the free parameters of the model can be defined as,

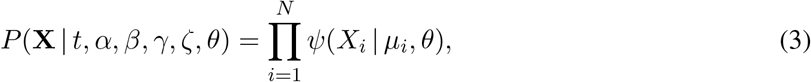

where *μ_i_* = *λP*(*Z_i_|t, α, β, γ, ζ*) and *P*(*Z_i_|t, α, β, γ, ζ*) is obtained from the matrix **P**(*t*) = *e*^**Q***t*^, as described above. As will be seen below, *λ* can generally be preestimated from global properties of a sequencing data set. Notice that typically there will be a data point corresponding to each state of the model, with the exception of state 0, which is unobservable.

### Nonequilibrium Case and Application to Time-Course Data

Suppose nascent RNA sequencing data have been collected in a time course, with read counts **X**^(*t*)^ for *t* ∈ {*t*_1_, *t*_2_, …, *t_M_*}. Typically *t*_1_ = 0 and represents the case prior to induction of transcription (Figure 4) [6]. Assuming independence of the data for the time points, a joint likelihood for all data can be defined as,

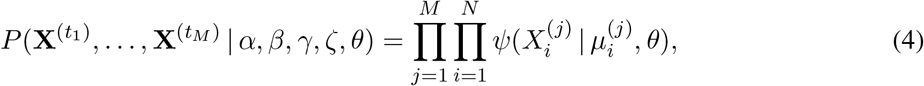

where 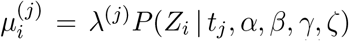, analogous to the case above. Notice that I allow here for a time-point-specific sequencing depth, *λ*^(*j*)^.

**Figure 4:**
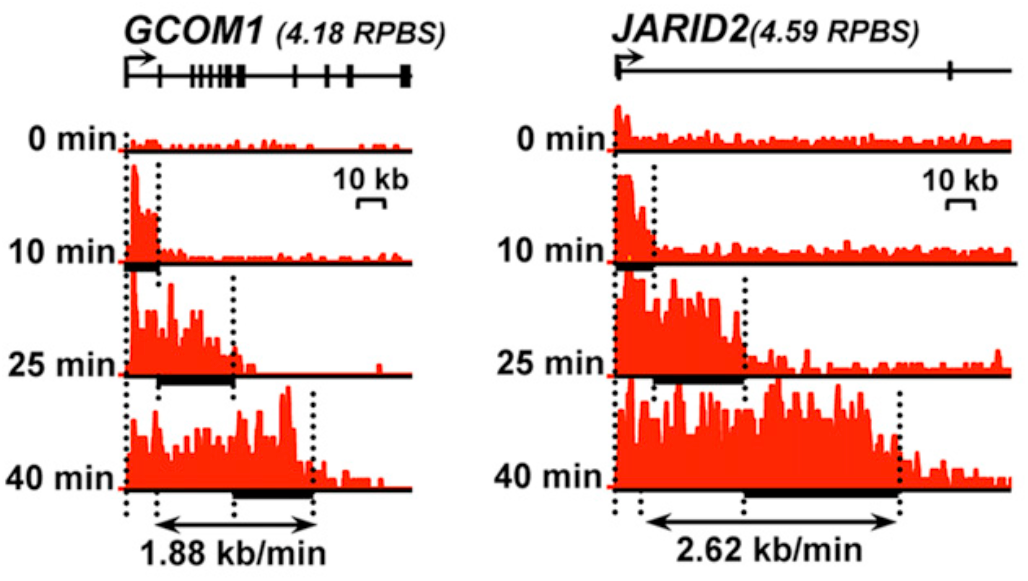
Examples of nascent RNA sequencing data collected in a time-course and used to estimate rates of elongation. Data were collected by GRO-seq at 0, 10, 25, and 40 minutes in MCF-7 cells [6]. Results are shown for the *GCOM1* (left) and *JARID2* (right) genes. In both cases, the left edges of the *x*-axes correspond to the annotated TSS. The RNAP “wave” can be seen to move rightward as time advances. Figure adapted from reference [6].

In this case, *α*, *β*, *γ*, and *ζ* can all be estimated from the data, at least in principle, although the information about some of them may be weak. However, there is a practical problem relating to polymerase positions that are impossible under the model (for example, when *t* = 0 and transcription has not yet commenced) but for which the corresponding read count *X_i_* is nonzero simply owing to the background noise of the assay. This problem can be addressed by defining *ψ*(*X_i_*|*μ_i_, θ*) by a suitable background distribution whenever *μ_i_* = 0 (or perhaps whenever *μ_i_ < T*, where *T* is a background-derived threshold). In this way, it should be possible to estimate an elongation rate *ζ* for each transcription unit directly from time-course data by numerical optimization of equation 4. This approach would provide an alternative to current hidden Markov model-based inference methods and would have the advantage of allowing estimation of *α* and *β* as well. The termination rate *γ* may be more difficult to estimate, but it may be possible to do so with a sufficiently long time course.

As observed above, *ζ* and *t* cannot be separately estimated from data, because they appear in the likelihood only as the product *ζt*. However, in a time-course setting, *t* is known and thus *ζ* can be estimated.

### Stationary Distribution and Inference at Steady-State

This Markov chain is ergodic and, therefore, will eventually reach a steady-state equilbrium defined by a unique stationary distribution over states. This stationary distribution, denoted ***π***, is invariant to *ζ* and can be found by solving the equation ***π***(**R**+ **I**) = ***π***, or equivalently, *π***R** = **0**. Owing to the simple structure of **R**, the solution for ***π*** can easily be determined by substitution.

Because state 0 is simply an abstraction to allow for the recirculation of RNAP and cannot be measured, we will ignore its stationary probability and instead describe the stationary distribution conditional on RNAP occupancy along the DNA template. This conditional stationary distribution is given by ***π*** = (*π*_1_, …, *π_N_*) such that,

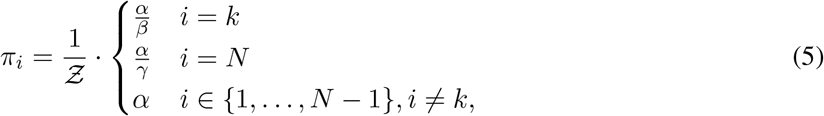

with normalization constant 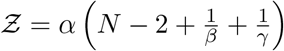.

This distribution has an intuitive interpretation. First, note that it will typically be the case that *β, γ* < 1, owing to slowdowns in elongation from pausing and termination. By contrast, RNAP should proceed unimpeded in the gene body, and therefore its density at steady state should reflect the initiation rate. Accordingly, *π_i_* is proportional to *α* in the gene body, but elevated by factors of 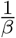 and 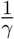 the pause peak and termination peak, respectively (Figure 5). The quantity 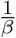 is sometimes estimated from nascent RNA sequencing data and described as the *pause index*. By analogy, the quantity 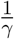 can be described as a *termination index*.

Notice that I have chosen to write the elements of ***π*** such that *α* appears in each numerator, despite that these *α* terms would cancel with the denominator and hence could be omitted. The reason for this choice will become clearer later in the article when multiple genes are considered, each with its own version of *α*. The inclusion of *α* in the stationary distribution will allow these values to be compared across genes.

**Figure 5:**
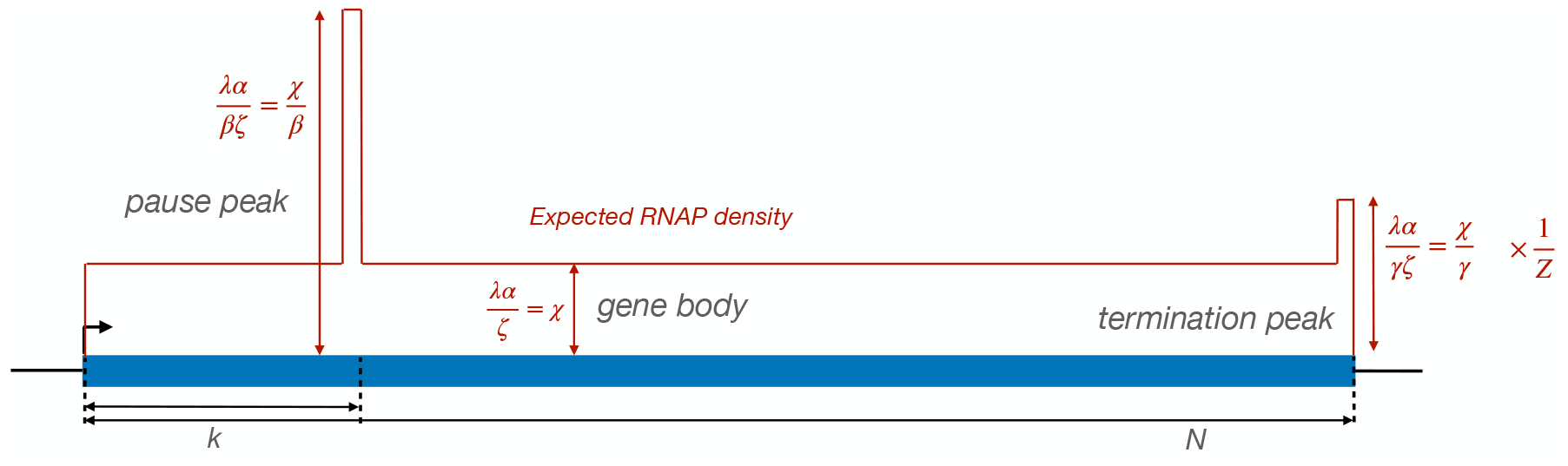
Expected density of RNAP at steady-state under the continuous-time Markov model.

Of course, when comparing different genes, it is possible that their elongation rates also differ. When examining nascent RNA-sequencing data at steady-state, the read depth in each TU *j* will be proportional to 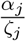, where *α_j_* and *ζ_j_* are, respectively, the initiation and elongation rates specific to that TU. Furthermore, the observed counts will be also be proportional to the sequencing depth, *λ*. Because these three parameters are not identifiable at steady state, I will represent them by the compound parameter 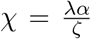. Thus, the stationary distribution can be written (Figure 5),

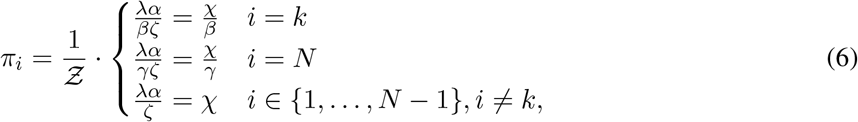

with 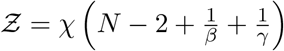.

This stationary distribution can be used to define a likelihood function for a system in steady state, described by a single data set (as opposed to a time course), denoted **X** = (*X*_1_, …, *X_N_*). In this case, let the expected read depth at each site *i* be *μ_i_ ∝ π_i_*, so that the likelihood of the data is given by (cf. equation 3),

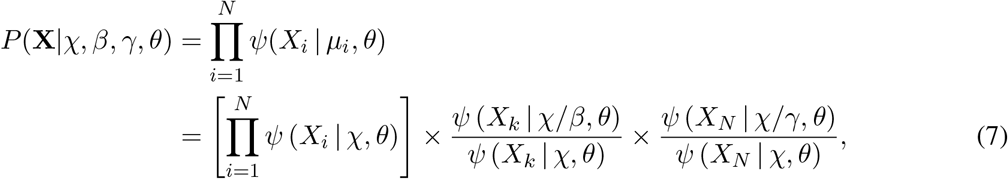

where the normalization constant for ***π*** is assumed to be absorbed by *χ*.

As it turns out, the termination peak is typically difficult to characterize with real nascent RNA sequencing data, owing to transcriptional run-on, poorly characterized 3′ ends of genes, and other factors. Therefore, from this point on, I will omit the termination component of the model and the corresponding parameter *γ*. With this simplification, the likelihood function can be written,

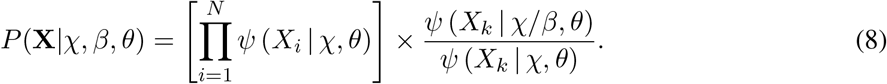

### The Poisson Case

In some applications, the generating distribution *ψ* may need to be complex to capture realistic patterns in sequencing reads, but in others, it may be sufficient to assume a simple form for this distribution. The assumption of a Poisson distribution at each nucleotide for *ψ* leads to particularly straightforward closed-form estimators in many applications of interest.

With a Poisson generating distribution, the steady-state likelihood becomes simply,

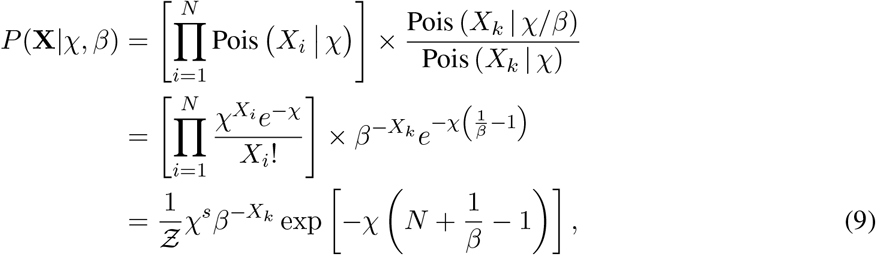

where 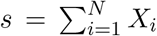 is a sufficient statistic equal to the sum of all read counts and 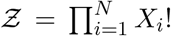 is a normalization term that does not depend on the model parameters and can be ignored during optimization. The log likelihood is given by,

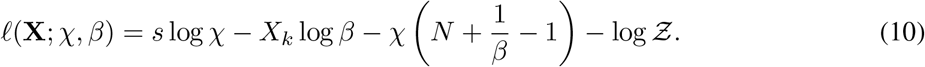

The maximum-likelihood estimators for *χ* and *β* have simple closed-form solutions:

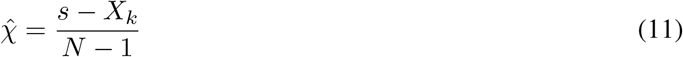

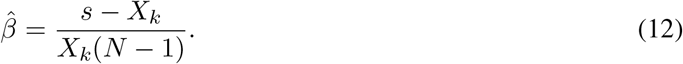

Thus, in this case, the estimator for *χ* is simply the average read depth, excluding the pause peak, and the estimator for *β* is the ratio of that same average read depth to the read depth in the pause peak. Notably, these are estimators that have been widely used in the analysis of nascent RNA sequencing data, with more heuristic justifications.

In practice, it is often best to avoid the complex signal in the pause region in estimating *χ* and instead estimate it from a downstream portion of the gene body. In this case, one can define 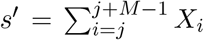,where *M* is the length of the interval considered, and estimate *χ* and *β* as,

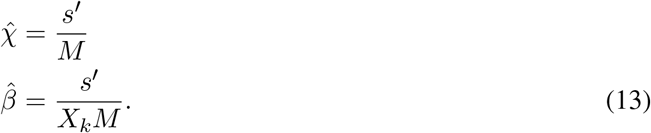

In most cases below, I will make use of these simpler, more robust estimators for *χ* and *β*.

While the nucleotide-specific counts in real data sets tend to be overdispersed, the simplicity and speed of the Poisson model may still make it an attractive option in many analyses. For example, as will be seen below, it may be desirable to perform statistical tests based on the Poisson model and then adjust *p*-values using an empirical calibration.

### Allowing for Variation in the Pause Site

In practice, the pause site tends to vary across cells, leading to a broad pause peak in real nascent RNA sequencing data. This problem can be addressed by allowing the location of the pause peak, *k*, to vary between some *k*_min_ and some *k*_max_ according to an appropriate distribution, and then assuming each read count *X_k_* within this range reflects a mixture of cells that do and do not have their pause site at position *k*. More specifically, I will assume that, for *k* ∈ {*k*_min_, …, *k*_max_}, *f_k_* represents the fraction of cells with pause site *k*, and that the read count *X_k_* derives from a mixture of one Poisson distribution with rate *χ/β · f_k_* and a second Poisson distribution with rate *χ* · (1 − *f_k_*). If we denote by *Y_k_* the (unknown) portion of the read count that derives from the first process, then the likelihood function can be expressed as (cf. equation 9),

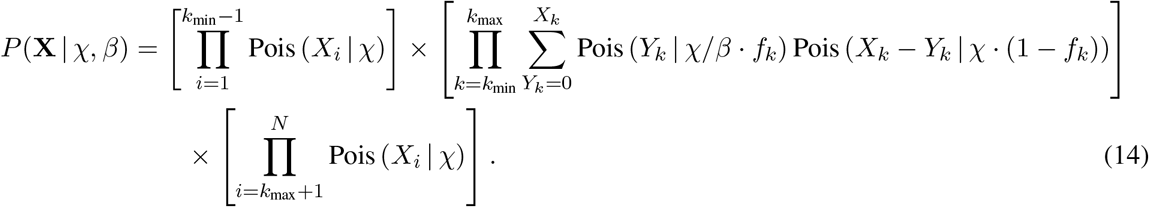

This likelihood function is straightforward to maximize by expectation maximization. The complete-data log likelihood function, with known values of *Y_k_*, can be expressed in terms of compact sufficient statistics (analogous to equation 10) as,

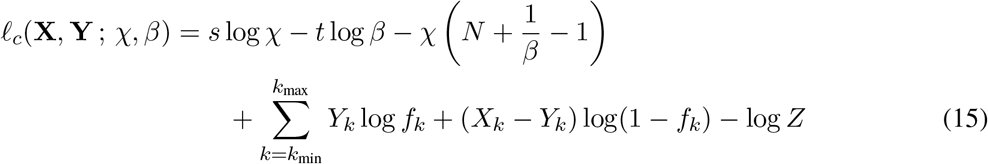

where *s* = ∑_*i*_ *X_i_* and *t* = ∑_*k*_ *Y_k_*. For simplicity, assume that *χ* is pre-estimated for a portion of the gene body downstream of *k*_max_ using equation 13. For the moment, assume also that the values of *f_k_* are fixed in a preprocessing step. Then *β* can be estimated as,

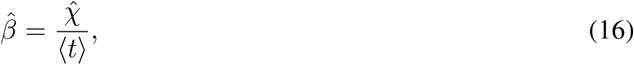

where 〈*t*〉 denotes the posterior expected value of *t*, which can be computed on each iteration by summing the posterior expected values 〈*Y_k_*〉. These values, in turn, can be computed by observing that, because *X_k_* is the sum of two Poisson-distributed variables, *Y_k_*|*X_k_* is binomially distributed with probability,

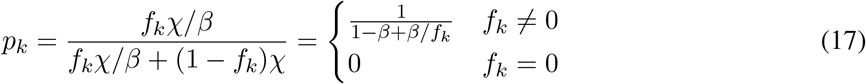

Therefore, 〈*Y_k_*〉 = *X_k_* · *p_k_*, and

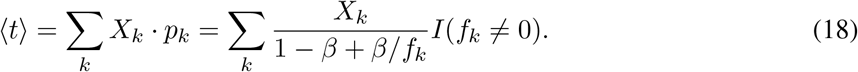

Thus, an EM algorithm can be implemented by iteratively applying equations 16 and 18, in turn, until convergence.

However, it may also be of interest to define the *f_k_* values in terms of a parametric distribution and estimate the parameters of that distribution also by EM. In this way, the distribution over pause sites can be estimated from the data at each TU of interest. A natural choice is a Gaussian distribution with mean *μ* and variance *σ*^2^,

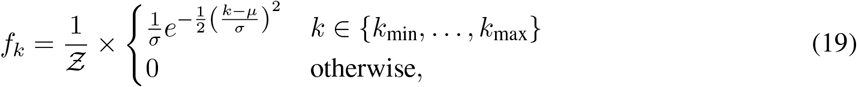

where the explicit normalization constant 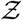 is needed because the distribution is applied to a bounded interval and is defined at integer values only.

In this case, it can be shown that the EM updates for *μ* and *σ*^2^ are simply,

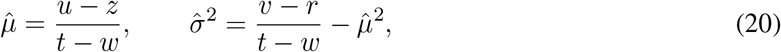

where,

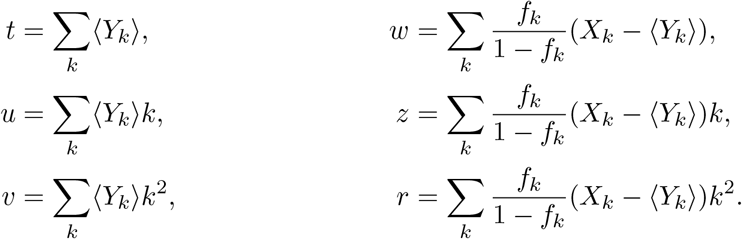

Notice that all of these quantities are easily computed from the 〈*Y_k_*〉 values together with *f_k_* and *X_k_*.

### Allowing for Collisions

As noted, the continuous-time Markov model described in this article makes the important simplification of ignoring collisions between polymerase molecules by modeling the movement of each RNAP independently of the others. In reality, an RNAP molecule can never pass another RNAP molecule along the DNA template, and will be blocked from forward progress if it catches up to its predecessor. The approximation should be adequate when the density of RNAPs per TU is low, but it will begin to fail when either the average density becomes high or when the elongation rate in certain local regions (such as the pause site) becomes sufficiently low that polymerases begin to back up.

#### Extension to the *ℓ*-TASEP

It turns out that the dynamics of “biopolymerization”—that is, the assembly of a protein or nucleic acid polymer by polymerases moving along a common nucleic acid template—has been investigated for decades (e.g., [18]). The relevant stochastic process is known in the literature as the *totally asymmetric simple exclusion process* (TASEP) [19]. In the case of heterogeneous elongation rates and a polymerase molecule that has an extended size *ℓ*—meaning that the centers of successive molecules can never be less than *ℓ* nucleotides apart—inference under this model is thought to be analytically intractable and is typically approached using Monte Carlo methods.

Interestingly, however, Erdmann-Pham et al. have recently given an approximate closed-form solution for the steady state density of this version of the model (the inhomogenous *ℓ*-TASEP) that considers pairs of interacting molecules [20]. Their solution was initially developed for the case of protein synthesis (i.e., with ribosomes moving along an RNA template), but in a subsequent work, they also applied it to the case of RNA transcription [21].

It appears that the solution of Erdmann-Pham et al. could be adapted in a fairly straightforward manner for the steady-state cases above. Essentially, one could discard the continuous-time Markov model and use the equations from ref. [20] in place of those given in equation 6. For example, this alternative stationary distribution could simply be substituted into equation 8 or equation 9 to obtain a new likelihood function. Statistical inference in this framework could be accomplished by numerical optimization. Notably, the generality of Erdmann-Pham et al.’s solution would allow it to be used also in the case where the elongation rate is different at each nucleotide position, as discussed later in the article.

Still, this solution to the *ℓ*-TASEP may be more complex than needed here. Instead, I will consider a simple extension of the Markov model introduced above that may be adequate for most analyses of eukaryotic transcription.

#### The special case of steric hindrance of initiation

In eukaryotic cells, it appears that, by far, the most pronounced reduction in elongation rate—and hence, the highest probability of collision—occurs at the promoter-proximal pause site. Indeed, based on current estimates of average rates of initiation and elongation, RNAP molecules should generally be fairly sparse along the gene body. For example, with an average elongation rate of 2.3 kb/min and an average initiation rate of 2.7 RNAP molecules per cell per minute [22], RNAPs would have an average spacing of 850 bp in the gene body. The rate of collision, of course, depends on other factors, such as the variance across cells in both initiation and elongation rate, as well as the variability over time in these rates (e.g., if transcription occurs in “bursts” collisions will be more likely), but it seems likely that, at least in many cases, collisions along the gene body will be infrequent enough that they can be ignored without too much cost.

Upstream of the pause site, however, collisions will be many times more likely. In addition, there is now considerable evidence that RNAP molecules frequently pile up behind the pause site to such an extent that they begin to block new RNAP molecules from initiating transcription. Because the pause site generally occurs ∼50 bp downstream of the TSS and the “footprint” of an engaged RNAP is ∼35 bp on the DNA template, the geometry seems to permit only about one, sometimes perhaps two or three, RNAP molecule to be held in the paused position before new initiation events are impeded [23]. This phenomenon could potentially lead to an effective decrease in the initiation rate owing to pausing—what Gressel et al. [22] have called the “pause-initiation limit.”

It turns out that the specific problem of a decrease in initiation owing to steric hindrance from paused RNAP molecules can be accommodated fairly easily in the framework introduced here, at least in the steadystate setting. If it is true that this is the dominant manner in which collisions impact nascent RNA sequencing data for eukaryotes, then the Markov model augmented to allow for steric hindrance may be adequate to describe many aspects of the complex interplay between initiation and pausing.

To define a simple model for steric hindrance, I will introduce a distinction between a *potential* rate of initiation in the absence of occlusion of the initiation site, *α*, and the *effective* rate of initiation after a portion of initiation events are blocked by an existing RNAP molecule, which I will denote *ω*. In particular, I will assume *ω* = (1 − *ϕ*)*α*, where *ϕ* is the probability of that the “landing pad” required for a new initiation event is already occupied by an RNAP. Thus, *ω* ≤ *α*. Notice that any estimation of initiation rates based on the density of RNAPs in the gene-body will be representative of *ω*, not *α*; a correction will be required to obtain an accurate estimate of the “true” rate *α*.

With a few mild simplifying assumptions, *ϕ* can be computed fairly easily. Assume first that the “footprint” of an engaged polymerase, *ℓ*, is sufficiently large that at most one RNAP can be present in this region at a time. (This assumption will be relaxed below.) Further assume that elongation up to position *k* occurs much faster than the initiation rate *αζ* or the pause-escape rate *βζ*, so that the dynamics of elongation through the pause peak can be ignored. In this case, occupancy of the landing pad can be described using a simple two-state continuous-time Markov model (Fig. 6A). Here, either the landing pad is unoccupied (state 0) and is therefore available for new initiation events, which occur at rate *αζ*; or the landing pad is already occupied by an RNAP (state 1) and no new initiation events can occur until that RNAP escapes from the pause site, which occurs at rate *βζ*. Thus, at steady state, the landing-pad occupancy *ϕ* is simply given by the stationary distribution of the occupied state,

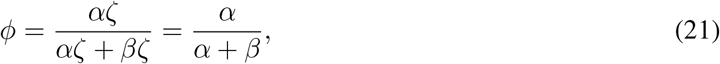

and the effective initiation rate, allowing for steric hindrance, is given by,

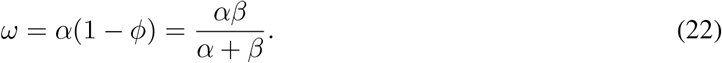

**Figure 6:**
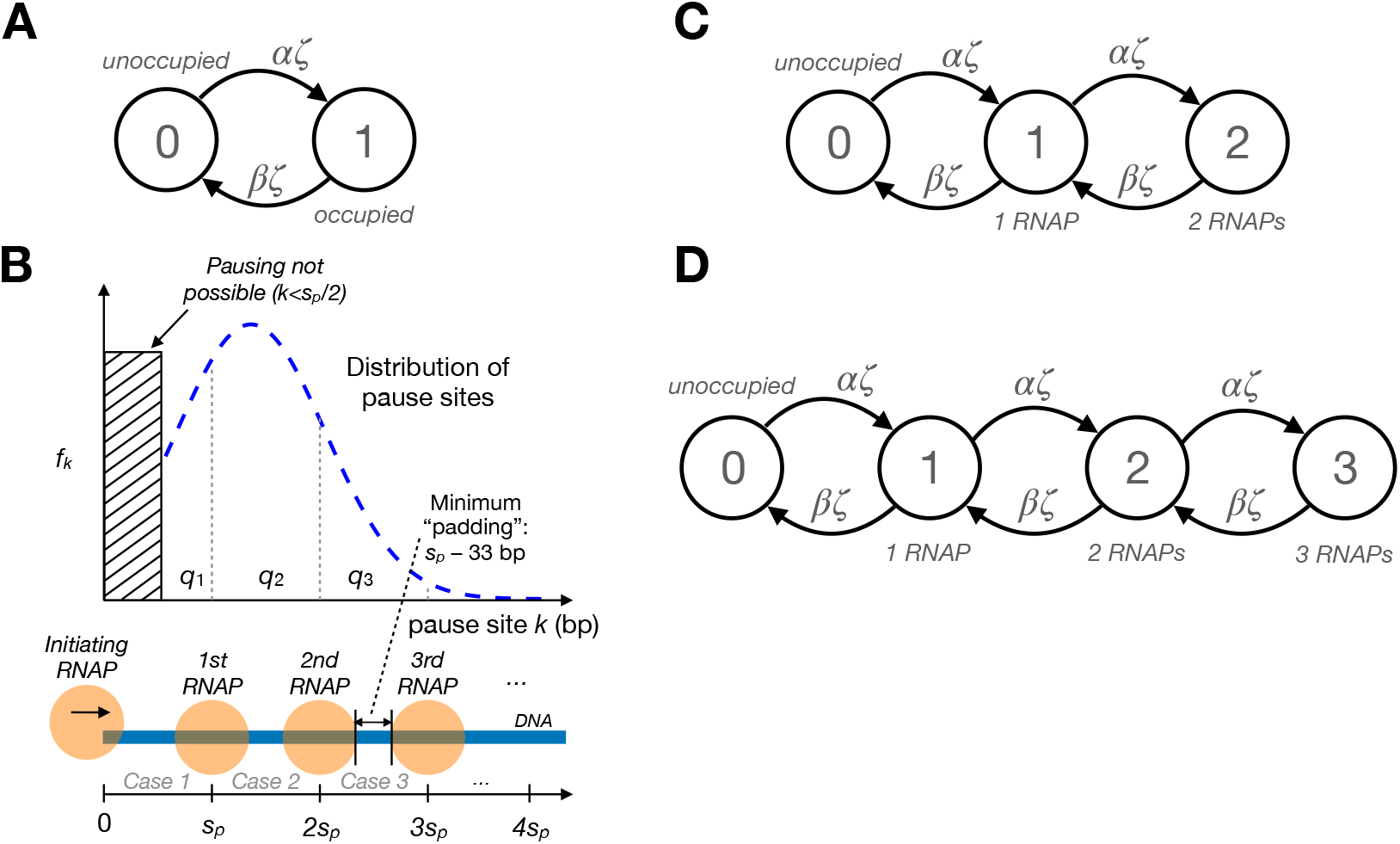
(A) Two-state continuous-time Markov model for steric hindrance of transcriptional initiation, assuming at most one RNAP at a time in the pause region. The pause region must be either unoccupied (state 0) or already occupied by another RNAP (state 1). Transitions from state 0 to state 1 occur at the (unimpeded) initiation rate, *αζ*, and transitions from state 1 to state 0 occur at the pause-escape rate, *βζ*. The stationary frequency of state 1 defines the landing-pad occupancy *ϕ* and is given by 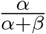. (B) Illustration showing a hypothetical distribution of pause sites *k* and its implications for the number of RNAPs that can simultaneously occupy the pause region. When *k≤s_P_*, where *s_P_* is the minimum center-to-center spacing between adjacent RNAPs, only one RNAP is possible (Case 1 in the text); when *s_P_ < k≤*2*s_P_*, up to two are possible (Case 2); and when 2*s_P_* < *k*≤3*s_P_*, up to three are possible (Case 3). Notice that the portion of the density corresponding to each Case *r* is given by *q_r_*. (C) Generalization of Markov model to accommodate up to two RNAPs in the pause region (Case 2). (D) Further generalization to accommodate up to three RNAPs (Case 3). The equation for *ϕ* can be generalized to account for these cases (see text).

Notice that these equations also imply that *ω* = *ϕβ*, meaning that, at steady state, the effective initiation rate *ω* must always be less than or equal to the pause-escape rate *β*, with *ω* approaching *β* as the landing-pad occupancy *ϕ* approaches unity. Therefore, if one estimates an effective initiation rate 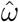 and a pause-escape rate 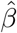 from the data, as described above, then one can obtain estimates of *ϕ* and *α* as follows,

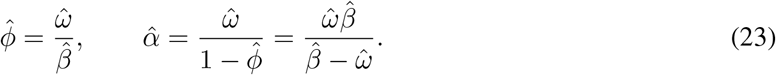

Notice that the estimator for *ϕ* will be proportional to the read-counts in the pause peak.

### Steric Hindrance with Multiple RNAPs

As it turns out, the assumption of ≤1 RNAPs per pause region is often too restrictive, and the presence of more than one RNAP in this region can have a substantial impact on the landing-pad occupancy *ϕ*. In this section, I generalize the model for steric hindrance to allow for any number *r* of RNAPs in the pause region, focusing in particular on the case of *r* ≤ 3, which should cover most plausible scenarios in human cells.

Let *s_P_* be the minimum center-to-center spacing, in nucleotides, between adjacent RNAPs on the DNA template. As noted in the main text, structural data suggests *s_P_* is at least 33 bp but more plausibly *s_P_*≈50 bp [23–26]. Further assume that a new initiation event can successfully occur if, and only if, the previous RNAP has advanced to a position *i > s_P_* (in other words, the “landing pad” for new initiation events has size *ℓ* = *s_P_*). Consequently, the maximum possible number of RNAPs in the pause region is *r* = 1 if the pause site *k* ≤ *s_P_*, *r* = 2 if *s_P_* < *k* ≤ 2*s_P_*, *r* = 3 if 2*s_P_* < *k* ≤ 3*s_P_*, and so on (see Fig. 6B). In general, 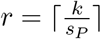.

In addition, assume a parobability mass function *f_k_* for the pause site *k* across cells, with cumulative distribution function 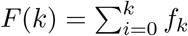. Let *q*_1_ be the *k* density associated with *r* = 1; that is, *q*_1_ = *F*(*k* = *s_P_*). Similarly, *q*_2_ is the density associated with *r* = 2, *q*_2_ = *F*(*k* = 2*s_P_*) − *F*(*k* = *s_P_*); and *q*_3_ = *F*(*k* = 3*s_P_*) − *F*(*k* = 2*s_P_*). In general, *q_r_* = *F*(*k* = *rs_P_*) − *F*(*k* = (*r* − 1)*s_P_*) (Fig. 6B).

Now, in each cell, *k* has a single value and therefore one of a series of mutually exclusive cases must apply. Let us denote by Case *r* (for *r* ∈ {1, 2, 3, *…*}) that the maximum possible number of RNAPs is *r*. Thus, in Case 1, *r* = 1 and *k* ≤ *s_P_*; in Case 2, *r* = 2 and *s_P_ < k* ≤ 2*s_P_*; and so on. Given the *f_k_* values, Case *r* must occur with probability *q_r_*. Therefore, *ϕ* can be calculated as a mixture of case-specific landing-pad probabilities, 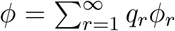, where *ϕ_r_* is the probability that the landing pad (first *s_P_* nucleotides) is occupied in case *r*. In practice, this quantity is approximated as 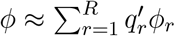, where *R* is the maximum plausible value of *r* (here, *R* = 3) and,

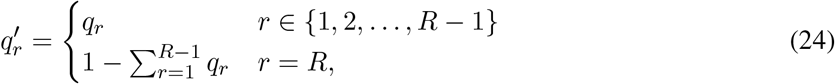

where the last term, 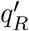, is “rounded up” to account for the remaining tail of the distribution.

The case of *r* = 1 has already been described in the previous section. It can be captured by a two-state model, where the landing-pad is either unoccupied (state 0) or occupied by a single RNAP (state 1; Fig. 6A). Therefore,

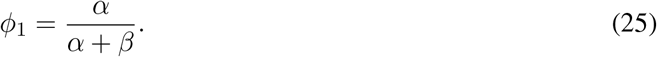

It turns out that the cases of larger values of *r* follow naturally through the addition of more states. For example, when *r* = 2, the landing pad is either unoccupied (state 0), occupied by one RNAP (state 1), or occupied by two RNAPs (state 2). Assuming no two events can occur simultaneously, state 2 can be reached only when a new initiation event occurs while state 1 is occupied (at rate *αζ*), and a pause-release event in state 2 causes a return to state 1 (at rate *βζ*). The result is a chain of states as shown in Fig. 6C. In addition, under the assumption that elongation through the pause peak is instantaneous, the landing pad is occupied if, and only if, state 2 is occupied. Thus, *ϕ*_2_ is given by the stationary frequency of state 2 in this model, which can be shown to be,

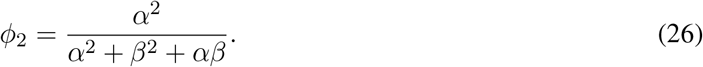

Similarly, the case of *r* = 3 can be addressed by extending the chain further with a state 3, and setting *ϕ*_3_ equal to the stationary frequency of that state (Fig. 6D),

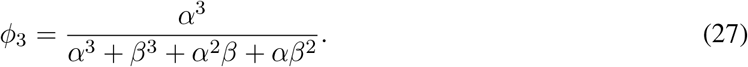

Therefore, assuming *R* = 3, *ϕ* can be estimated as,

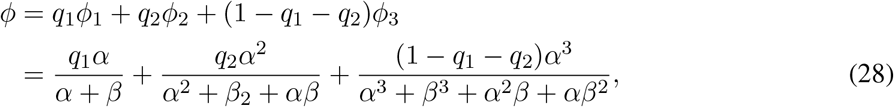

allowing for the possibilities of one, two, or three RNAPs in the pause region of each cell. As *s_P_* grows larger and/or *f_k_* shifts toward the TSS, the fraction *q*_1_ approaches 1, and equation 28 approaches equation 21.

As it turns out, substitution of *ω/*(1 − *ϕ*) in place of *α* in this more general expression for *ϕ* leads to a complex polynomial that cannot be easily solved for *ω*. Instead, this equation is solved numerically to obtain *ϕ* and *α* from *ω* and *β* (analogous to equation 23).

#### Fitting the Steric-Hindrance Model to Data

The multi-RNAP steric hindrance model can be combined with the model for variable pause sites across cells and fitted to the data by a relatively straightforward extension of the EM algorithm described above. As in that case, assume that the unscaled initiation rate, *χ*, is pre-estimated from data in the gene body. In addition, however, this case requires pre-estimation of the scale-factor *λ* and the elongation rate *ζ*, so that a scaled estimate of the effective initiation rate, 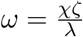, can be obtained from each estimate of *χ*.

In addition, we must address the problem that the landing-pad occupancy *ϕ* is constrained to fall between 0 and 1, but the relationships above do not enforce such a constraint. For example, in the single-RNAP case, where 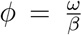 (equation 23), *ϕ* will be undefined whenever *ω > β*. A similar (but more complicated) relationship holds in the multi-RNAP case.

To address this problem, I take a Bayesian approach and assume a (weakly) informative prior distribution for *ϕ*. This strategy not only restricts *ϕ* to the allowable range but has the benefit of regularizing the model when information in the data about *ϕ* is weak. With the assumption of a Beta(*ϕ|a, b*) prior for *ϕ*, with shape parameters *a* and *b*, the likelihood becomes,

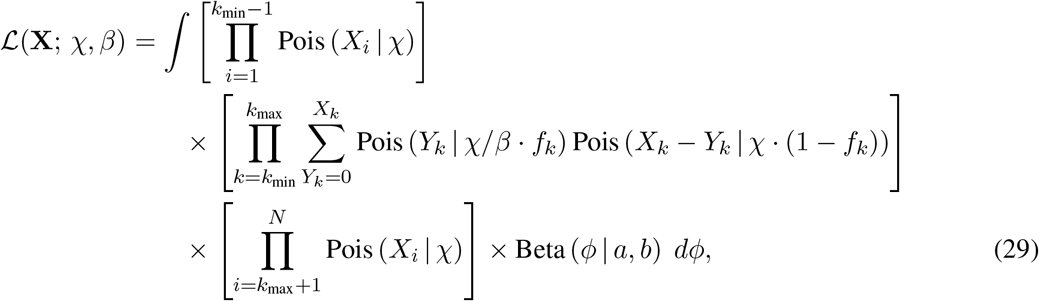

and the expected complete-data log likelihood (cf. equation 15) becomes,

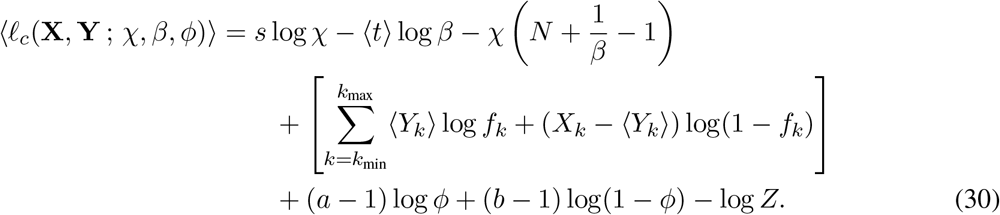

From a comparison of equations 15 and 30, it is evident that the calculation of the summary statistics 〈*t*〉, 〈*u*〉, 〈*v*〉, 〈*w*〉, 〈*z*〉, and 〈*r*〉 and the updates for *μ* and *σ*^2^ will all remain unchanged. However, this simplified presentation obscures that the parameters *ϕ* and *β* are implicitly linked by a function, *ϕ* = *g*(*β*; *χ*), which is indirectly defined by equation 28 as well as by the relationship *ω* = (1 − *ϕ*)*α*. Thus, the update for *β* in this case is no longer that shown in equation 16 but now must also consider the terms that depend on *ϕ*. As it turns out, in the full multi-RNAP model, rewriting equation 30 in terms of *β* only (i.e., by substitution for *ϕ*) leads to rather unwieldy polynomial expressions. Nevertheless, the M-step in the EM algorithm can be performed numerically without much trouble. In particular, on each iteration of the algorithm, all expected sufficient statistics are computed as before (E-step), but then, for the M-step, *β* is estimated by numerically maximizing the portion of equation 30 that depends on *β* and *ϕ*, that is,

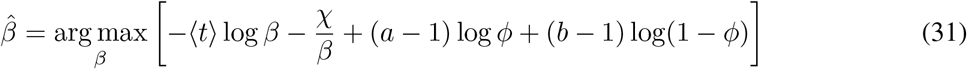

subject to the constraints of equation 28 and the relationship *ω* = (1 − *ϕ*)*α*. Thus, the previous EM algorithm can be adapted to the multi-RNAP steric hindrance case by simply replacing the M-step for *β* with this numerical optimization. No other changes are required.

### Allowing for Overdispersion

In general, read counts in nascent RNA sequencing data tend to be overdispersed (with variance exceeding the mean), just as in RNA-seq data. A relatively straightforward way to allow for overdispersion is to assume a mixture of Poisson rates within each region while maintaining the assumption of independence across nucleotide sites. In this way, some sites are allowed to accumulate reads at higher-than-average rates, and others at lower-than-average rates. If the mixture over rates is assumed to be Gamma-distributed, then this assumption simply implies that the site-specific Poisson distributions for read counts are replaced with negative binomial distributions.

In particular, if the read depth at each nucleotide *i* is assumed to be scaled by a random variable *ρ_i_*, which is Gamma-distributed with shape parameter *A* and rate parameter *A* (so that *E*[*ρ_i_*] = 1), and the assumption of independent Poisson read counts is maintained as in equation 9, then the likelihood function becomes,

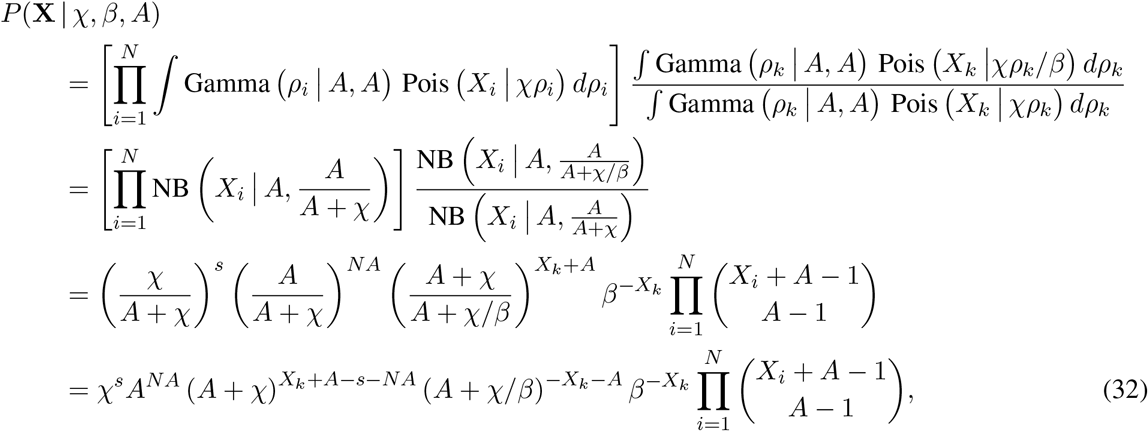

where NB(*X|r, p*) indicates that *X* has a negative binomial distribution with parameters *r* and *p*, that is, such that,

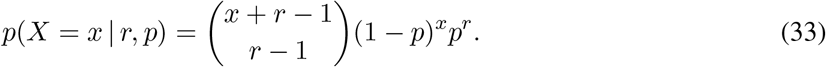

Notice that *A* must be a positive integer for this likelihood function to be well-defined.

In general, there are no convenient closed-form expressions for the maximum likelihood estimates of the parameters in this case, and they would need to be estimated numerically or by expectation maximization. For this reason, I will focus on the Poisson model in many of the sections below (but see Discussion).

### Likelihood Ratio Tests for Differences Between Transcription Units, Conditions, or Species

These likelihood functions lead naturally to a class of likelihood ratio tests (LRTs) for comparing aspects of the transcriptional dynamics of different TUs, or of the same TU under different conditions. For example, one might test whether two genes have different initiation rates or different pause-release rates. In each case, it might be desirable to allow some parameters to vary freely, while testing for evidence in the data of a difference in another parameter. For example, one might test for a difference in the pause-release rate allowing for differences in the initiation rate, or a difference in the initiation rate allowing for differences in the pause-release rate. I will focus on tests based on steady-state data, but one could equally well conduct a test for a difference in elongation rate based on time courses representing different conditions.

These LRTs all have the same general form. Consider read counts captured at steady-state for two TUs, 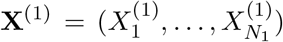 and 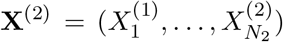. As noted, **X**^(1)^ and **X**^(2)^ could represent the same TU under different conditions, different TUs under the same conditions, or even different TUs under different conditions. In a comparative genomic analysis, they may represent orthologous TUs in different species (say, human and chimpanzee). Let the free parameters for the two genes be denoted Θ_1_ = (*α*_1_, *β*_1_, *θ*_1_) and Θ_2_ = (*α*_2_, *β*_2_, *θ*_2_), respectively (ignoring the *ζ* parameters for now, and focusing on the steady-state), where *θ*_1_ and *θ*_2_ represent any additional parameters of interest. Finally, let the log likelihood of a particular data set be denoted *ℓ*(*X*; Θ), and let the maximized value of that likelihood be denoted 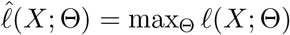.

In general, one wishes to compare a null hypothesis *H*_0_ that the two data sets can be jointly described by (at least some) shared parameters against an alternative hypothesis *H_A_* that they require different parameters. The parameter can be partitioned into three mutually exclusive and exhaustive categories:

- *Fully tied* parameters (Θ_T_). These parameters are shared by both TUs under the null and alternative hypotheses.
- *Fully free* parameters (Θ_F1_ and Θ_F2_). These parameters vary freely in the two data sets under both the null and alternative hypotheses.
- *Tested* parameters (Θ_*_ or Θ_*1_ and Θ_*2_). These parameters vary freely under the alternative hypothesis but are tied under the null hypothesis.

With these assumptions, the test statistic for an LRT for a difference in the tested parameters can be defined generally as,

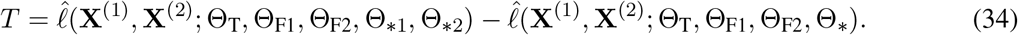

Here, each term represents the joint likelihood of the two data sets. Generally, **X**^(1)^ and **X**^(2)^ will be assumed to be conditionally independent given the parameters, so that their joint likelihood is a product of their individual likelihoods; however, I present them in this more general form to emphasize the need to estimate tied parameters jointly. In the usual manner, the first term represents the maximized likelihood under the alternative hypothesis and the second term represents the maximized likelihood under the null hypothesis. If the tested parameters, Θ_*1_ and Θ_*2_, are completely unconstrainted under the alternative hypothesis, *T* represents a two-sided test. However, they can be constrained, if desired, to achieve a one-sided test (for example, allowing only *α*_2_ ≥ *α*_1_ to test for an increased initiation rate in the second TU).

To make this framework concrete, let us consider some specific examples of tests. First, suppose one wishes to test for a difference in initiation rate between two TUs from the same data set, assuming equal elongation rates. In this case, Θ_T_ = {*λ, ζ*}, Θ_F1_ = {*β*_1_}, Θ_F2_ = {*β*_2_}, and Θ_*_ = {*α*}.

Next, suppose one wishes to test for a difference in the pause-release rate between two TUs from the same data set. In this case, Θ_T_ = {*λ*}, Θ_F1_ = {*α*_1_, *ζ*_1_}, Θ_F2_ = {*α*_2_, *ζ*_2_}, and Θ_*_ = {*β*}.

Finally, suppose one wishes to test for a difference in initiation rate in the same TU under two conditions, represented by separately sequenced nascent RNA sequencing data sets. Here, Θ_T_ = {*ζ*}, Θ_F1_ = {*β*_1_, *λ*_1_}, Θ_F2_ = {*β*_2_, *λ*_2_}, and Θ_*_ = {*α*}.

In general, for two-sided tests, the asympotic distribution of 2*T* under the null hypothesis will be a chisquare distribution with |Θ_*_| degrees of freedom, enabling convenient calculation of nominal *p*-values. In many cases, however, it may be preferable to use empirical *p*-values instead, for example, by computing *T* for a collection of TUs believed to be representative of the null hypothesis, and using this empirical distribution in place of the null.

### Poisson-based Likelihood Ratio Tests

In the case of the Poisson generating distribution at steady state, it is often possible to find closed-form solutions for the test statistic *T*. For example, in the test for differences in initiation rate (*H*_0_ : *α*_1_ = *α*_2_ vs. *H_A_* : *α*_1_ ≠ *α*_2_), *T* can be expressed as,

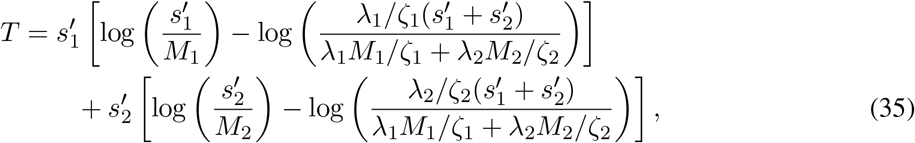

where the simplified estimation method in equation 13 is assumed, and where 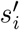 denotes the sum of read counts and *M_i_* denotes the length of the gene body in TU *i*. Thus, the LRT for a difference in *α* depends only on the data in the gene body, not on the pause peak.

It is sometimes convenient to re-express *T* in terms of weights *τ*_1_ and *τ*_2_, such that,

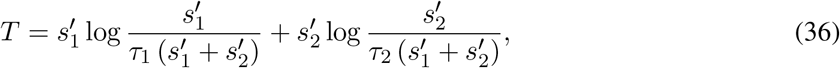

where,

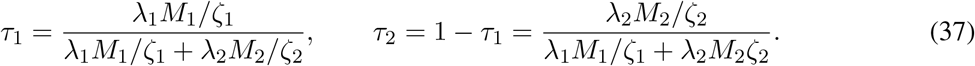

When *λ*_1_ = *λ*_2_, as when comparing the read counts for two different TUs from the same data set, when *M*_1_ = *M*_2_, as can typically be ensured by construction, and assuming *ζ*_1_ = *ζ*_2_, the weights are equal 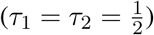 and *T* reduces to an expression that depends only on 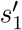 and 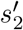:

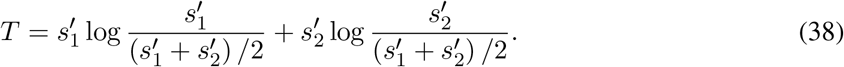

In this way, *T* can be seen to be a measure of discordance in read depth, normalized for differences in library size, gene-body length, and elongation rate. Notice in particular that, when the (normalized) average read depths are the same in the two gene bodies (e.g., when 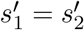 in equation 38), then *T* = 0 and the null hypothesis cannot be rejected. *T* grows larger as these average read depths become more different from one another. Importantly, however, *T* depends on the raw read counts, not only on their ratios. Thus, *T* tends to be small when the 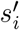 values are small, and to grow larger as they increase, reflecting greater confidence in the rejection of the null hypothesis with more data.

The test statistic for a difference in pause-release rates *β*_1_ and *β*_2_, with a fixed pause site *k* but allowing for a difference in initiation rates, is messier but can still be expressed in closed form as,

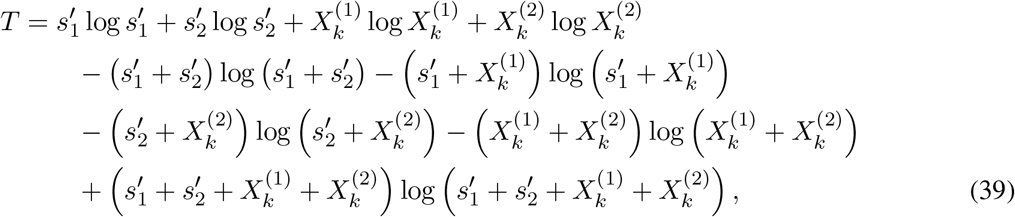

where, for simplicity, I focus on the case of shared *M*_1_ = *M*_2_ = *M* and further assume *k*_1_ = *k*_2_ = *k* (this test is independent of *λ*_1_, *λ*_2_, *ζ*_1_, and *ζ*_2_). Here, it can be shown easily that *T* = 0 if the ratios of read counts at the pause peak and the gene body are equal, i.e., if 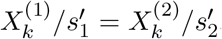, and that *T* grows larger as these ratios become more divergent, but in a manner that depends not only on the ratios but also on the absolute counts. Notably, this test can be shown to be a special case of a *G*-test and asymptotically equivalent to both a chi-squared test and Fisher’s exact test based on the four counts, 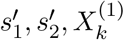 and 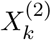.

The test for a difference in *β* becomes more complicated when allowing for uncertainty in the pause site, but this situation be accommodated relatively easily by taking advantage of the EM framework and precomputed posterior expected values of *t*^(1)^ and *t*^(2)^, which can be used in place of 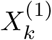 and 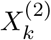, respectively. In particular, the test statistic is exactly the same as in equation 39 but with these posterior expected values in place of observed ones,

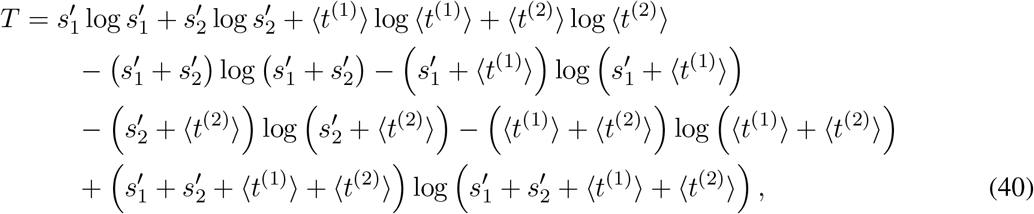

where the quantities of the form 〈*t*^(*j*)^〉 are computed iteratively as in equation 18. Strictly speaking, however, the 〈*t*^(*j*)^〉 values that appear in combination with other parameters, which reflect *H* _0_ (lines 2–4 of equation 40) should be calculated using the version of the model where *β* is tied, and the 〈*t*^(*j*)^〉 values that appear alone, which reflect *H _A_* (line 1), should be calculated using the version of the model with free parameters *β*_1_ and *β*_2_. The EM algorithm for the tied parameters is similar to the one for free parameters (equations 16–18) but the update for the shared *β* on each iteration is,

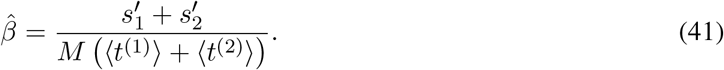

The iterative calculation of 〈*t*^(1)^〉 and 〈*t*^(2)^〉 is as specified in equation 18 but using the shared value of *β*.

Finally, the LRT for a difference in initiation rate can be extended to allow for steric hindrance. By explicitly allowing for interactions between pausing and initiation, it is possible to test whether an observed difference in initiation rates is beyond what can be explained by steric hindrance alone. In other words, this more conservative test excludes the possibility that what appears to be a difference in initiation rates is simply a consequence of different effects of the pause-initiation limit.

In this case, we assume *ω*_1_ = *α*_1_(1 − *ϕ*_1_) and *ω*_2_ = *α*_2_(1 − *ϕ*_2_) such that *α*_1_ = *α*_2_ under the null hypothesis and *α*_1_ ≠ *α*_2_ under the alternative hypothesis. As above, *β*_1_ and *β*_2_ are free parameters. As it turns out, the test statistic *T* is equivalent to the one given in equation 38, but with the weights *τ*_1_ and *τ*_2_ redefined to incorporate the terms (1 − *ϕ*_1_) and (1 − *ϕ*_2_):

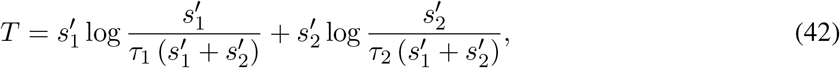

where,

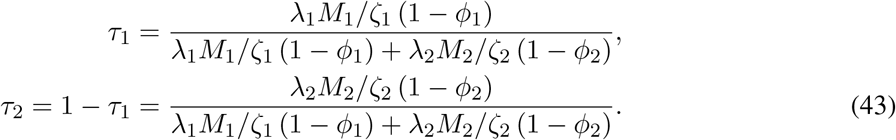

In the case where *k* is unknown, the test can be executed by fitting the model by EM under the null and alternative hypotheses.

### Bayesian Inference Under the Poisson Model

Another advantage of the Poisson model at steady-state is that it leads to relatively straightforward Bayesian inference of the key model parameters, *χ*, *β*, and *γ*. To simplify the mathematics, I will introduce a change of variables from *β* to 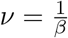 and assume Gamma priors for *χ* and *ν*:

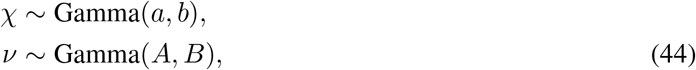

where the first parameter of each Gamma distribution represents its shape and the second parameter represents its rate (inverse scale).

With these assumptions, and assuming *k* is known, the joint posterior distribution of the parameters is given, up to a normalization constant, by:

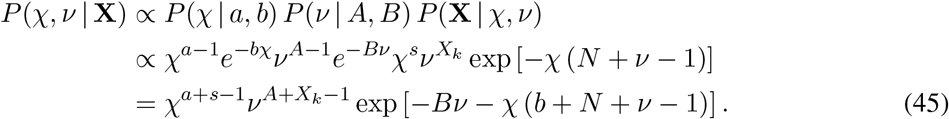

Because *χ* is entangled with *ν* in the last term of equation 45, the marginal posterior distributions for the individual parameters do not have standard forms. However, one can sample from the joint posterior distribution by rejection sampling or sampling-importance resampling using the unnormalized density in equation 45.

In addition, it can be shown that the conditional distribution of each parameter given the others reduces to a standard Gamma distribution:

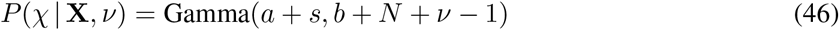

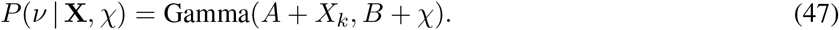

As a result, it is straightforward to implement a Gibbs sampling algorithm to sample from the joint posterior, from which marginal posterior distributions can easily be approximated. Using this framework, it would be possible to calculate Bayes factors for model comparison, analogous to the LRTs introduced above.

In the case of the steric-hindrance model, the posterior distributions of *ω* and *ϕ* can be obtained from those for *χ* and 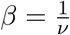 in post-processing, using equation 21.

### Generalization to Site-Specific Elongation Rates

The model can easily be generalized to allow for a different elongation rate *ζ_i_* at each nucleotide position *i*, instead of a constant rate across all nucleotides. In this case, the elongation rate no longer serves as a simple scaling factor for the infinitesimal generator matrix **Q**, and there is potentially information in the data about site-specific elongation rates, even at steady-state.

In this case, I will ignore the effects of both pausing end termination, and focus on the gene body, where the elongation signal is easiest to interpret. Thus, with site-specific elongation rates *ζ_i_*, the conditional stationary distribution for *i* ∈ {1, …, *N*} becomes simply (cf. equation 6):

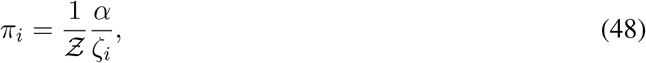

with normalization constant, 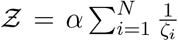, where I now assume that all positions *i* ∈ {1, …, *N*} fall in the gene body. Notice that I have maintained the parameter *α* here, despite that it appears only as a compound parameter together with the site-specific elongation rates and hence is nonidentifiable. The reason will become clear in the next section.

### A Generalized Linear Model for Discovering Features Associated with Elongation

In practice, nascent RNA sequencing data tends to be quite noisy at individual nucleotides, and it is not likely to be useful for direct inference of each *ζ_i_* value. However, it may be possible to learn something about the site-specific elongation process by pooling information across many sites and many TUs.

In particular, I will define a generalized linear model to make use of a collection of generic features at each transcribed nucleotide across the genome. These could include any property known or suspected to influence elongation r ates, i ncluding, for example, the degree of chromatin accessibility in a cell type of interest (assayed by DNase-seq or ATAC-seq), proximity to a splice site, presence of nucleosomes (MNaseseq), presence of stem-loops or other secondary structures of RNAs, or local G+C composition. Suppose there are *D* such features, summarized by a vector **Y**_*i*_ at each site *i*. Let *ζ_i_* be an exponentiated linear function of these features,

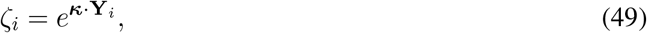

where ***κ*** is a vector of *D* real-valued coefficients shared across all sites, and the function *e ^x^* ensures that *ζ_i_* takes a positive value. It may be useful to define the first element of **Y**_*i*_ to be a constant of 1 to accommodate an intercept for the linear function at the corresponding position in ***κ*** (implying *D* − 1 true features).

In order to separate global aspects of the elongation process from TU-specific (and highly variable) rates of initiation, I will maintain a separate instance of the parameter *α* for each TU, but share the coefficient vector *κ* across all TUs. Thus, for *M* (independent) TUs indexed by *j*, the joint log likelihood function under the Poisson model is (cf. equations 9 & 10),

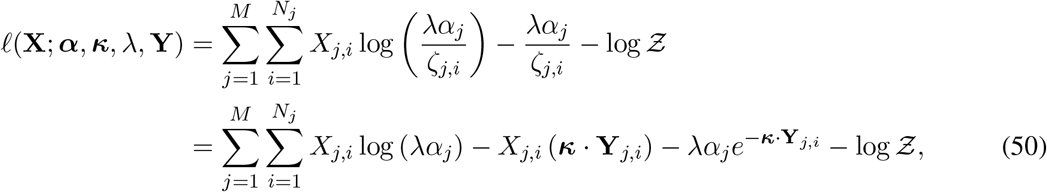

where *α_j_* is the initiation rate and *N_j_* is the total length for TU *j*; and *X_j,i_* is the read-count, **Y**_*j,i*_ is the feature vector, and *ζ_j,i_* is the elongation rate at the *i*th nucleotide in TU *j*;

#### Fitting the GLM to data

The log likelihood can be written in terms of simple summaries of the data, as follows,

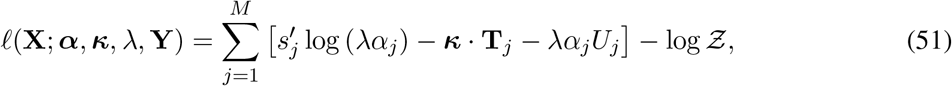

where 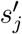 is the now-familiar sum-of-read-counts for the gene body, but restricted here to the *j*th TU, and **T**_*j*_ and *U_j_* are analogously defined as,

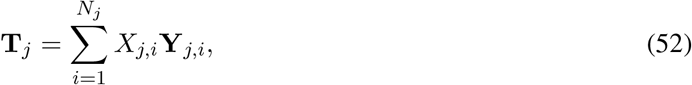

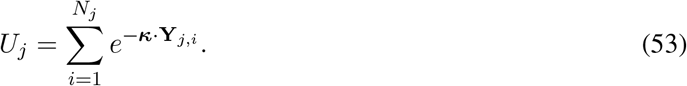

Notice that *U_j_* is not strictly a sufficient statistic because it depends on the parameter ***κ*** as well as on the data.

This joint log likelihood can be maximized by gradient ascent. The partial derivative with respect to the *n*th component of ***κ*** is given by,

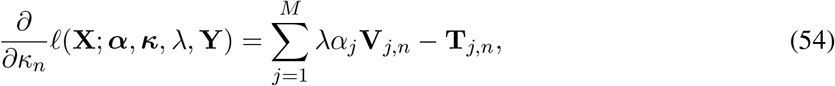

where the final subscript *n* indicates the *n*th element of a vector, and **V**_*j*_ is defined as,

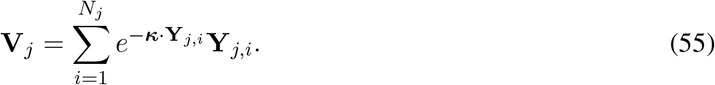

For a given value of ***κ***, the maximum for *α_j_* can be determined analytically as,

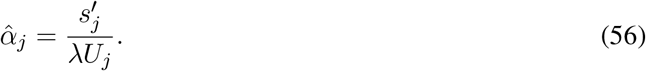

Thus, a gradient ascent algorithm can be implemented to iteratively improve estimates of ***κ*** and, on each iteration, to fully optimize *α_j_* conditional on the previous parameter values. Importantly, the sufficient statistics 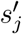 and **T**_*j*_ need only be computed once, in pre-processing, although *U_j_* and **V**_*j*_ must be recomputed on each iteration of the algorithm.

Notably, these results are closely related to those for standard Poisson regression. As in that case, the log likelihood function is guaranteed to be convex.

#### Filters for genomic features

In applications to real data, the genomic features will generally require some preprocessing, because they will have different noise properties and levels of resolution along the genome. A general and flexible way to convert them to a common representation that is matched to the nascent RNA sequencing data is to apply a separate smoothing filter to each feature.

In particular, let a filter 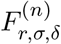 for each feature *n* ∈ {1, …, *D*} be defined by three parameters: a *radius* of application *r*, a *smoothing bandwidth σ*, and an *offset δ* (which allows for a translation along the genome sequence in the relationship between the covariate and the data *X_i_*). Each (scalar) covariate 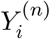 can then be replaced with a filtered version,

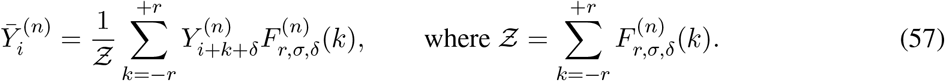

The filter 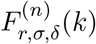 can take a variety of forms. For example,

- Gaussian filter: 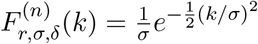
- Laplace filter: 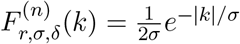
- Negative exponential filter: 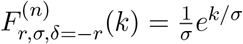
- Positive exponential filter: 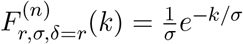
- Sine filter: 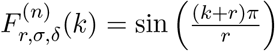.
- Cosine filter: 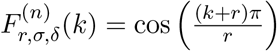.
- Averaging filter: 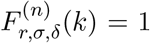. The *identity* filter 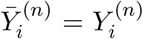 can be obtained by the special case, 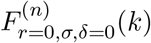.
- Generalized filter: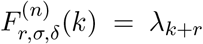, which is defined by a vector of nonnegative scale factors, ***λ*** = {*λ*_0_, *…, λ*_2*r*_}.

Notably, these filters can also be *composed* to define a richer, more general family of filters. For example, a composition filter 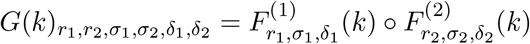 can be defined such that,

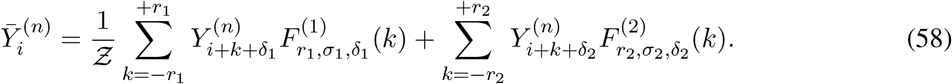

In principle, the parameters that define the filters could be estimated from the data along with the parameters of the GLM, but this approach is likely to be unwieldy and computationally expensive. Instead, it will most likely be preferable to determine them in a preprocessing step by examining each of the covariates in turn. For example, it may be possible to arrive at reasonable choices of a filter and its parameters by examining metaplots of nascent RNA sequencing readcounts centered on features of interest, such as ChIP-seq or ATAC-seq peak calls, or splice sites called from RNA-seq data.

### Technicalities

#### Estimation of *λ* and interpretation of rate parameters

As noted, the parameter for sequencing depth, *λ* is confounded with those for TU-specific initiation and elongation rates, *α_j_* and *ζ_j_* at steady-state, in the sense that all three parameters appear only in the relevant likelihood functions as the product 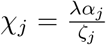. Because these parameters all contribute linearly to increases in the expected read counts, they cannot be independently estimated without some exogenous calibration, which in most cases will not exist.

The most pragmatic path forward seems to be to estimate *λ* in a preprocessing step, hold it fixed throughout the analysis, and interpret estimates of *α_j_/ζ_j_* relative to each other and to the meaning of *λ*. Notably, *β_j_* must be interpreted in the same relative manner, since it is effectively tied to *α_j_* and *ζ_j_* through the structure of the continuous-time Markov model. In some cases, it may be reasonable to assume *ζ_j_* is constant across TUs (as when comparing orthologs between closely related species), in which case relative values of *α_j_* can be obtained. Overall, it is worth emphasizing that the general framework presented in this article should be regarded as a means for estimating *relative*, rather than absolute, rates of initiation, pause-release, and, potentially, elongation. Nevertheless, these rates can effectively be compared with each other, across TUs, and—with appropriate care, as discussed below—across experiments.

Two possible ways to estimate *λ* are to make use of (1) the average read counts across all TUs, or (2) the read counts at one or more designated “reference” TUs. In either case, it would be sensible to make use of the gene bodies only of confidently annotated and likely expressed TUs. If the set of such TUs is denoted *V* (where *V* is a large, inclusive set in the first case or a smaller, more restricted one in the second case), then *λ* can be estimated in preprocessing simply as,

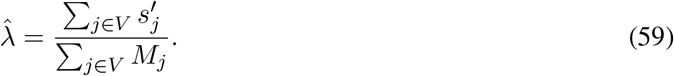

(Notice that this formula will implicitly weight TUs by length; if desired, an unweighted average of averages could be used instead.) After estimation of 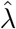, an initiation/elongation rate ratio can be determined for each TU *j* as,

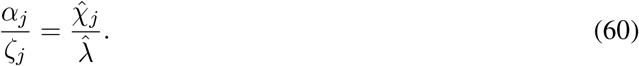

The choice of *V* will determine the meaning of these *α_j_ /ζ_j_* ratios, and in turn, the meaning of the *β_j_* parameters. If *V* is taken to include all (expressed) TUs, then *α_j_ /ζ_j_* will be expressed relative to the average behavior across the genome, with *α_j_ /ζ_j_* = 1 indicating an average initiation/elongation rate ratio, *α_j_ /ζ_j_* < 1 indicating a lower-than-average such ratio, and *α_j_ /ζ_j_ >* 1 indicating a higher-than-average such ratio. If instead, *V* is a set of unusually highly or lowly expressed genes, then *α_j_ /ζ_j_* must be interpreted accordingly.

It is worth noting that if one were to have access to a set *V* of TUs with known absolute values of *α_j_* and *ζ_j_*, then these TUs could serve as a calibration for the *α_j_ /ζ_j_* ratios, allowing them to be be assigned absolute, rather than relative, values.

#### Batch effects and normalization

A major practical problem in the analysis of all transcriptomic data, including nascent RNA sequencing data, is bulk differences in the distributions of values from separate experiments that are not biologically meaningful but instead reflect differences in sample or library preparation, sequencing, or some other technical aspect of data collection. In our case, such “batch effects” could be particularly problematic in tests of differences in initiation or pause-release rates across different conditions or species. If they are ignored, they may produce many false-positive predictions of differences.

In many cases, batch effects can be effectively addressed in a preprocessing step that eliminates systematic differences across many TUs that are likely to reflect technical artifacts, leaving differences more likely to be driven by the biological processes under study. I will focus here on quantile normalization [27], although the discussion will apply to other normalization methods as well, such as ones based on principal component analysis.

Assume that quantile normalization is applied to the gene bodies of all TUs in each data set under study, such that the original value of 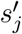 is replaced by a new normalized value 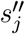. For now, assume that a proportional correction is made to the read counts across the TUs (including in the pause and termination peaks), so that each count *X_j,i_* is replaced by 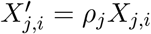 where 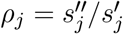.

One possible strategy would be to apply the methods described in this paper directly to the normalized data set, without change. An issue with this approach is that the normalized read counts, in general, will no longer have integer values, and for some applications, they would either need to be rounded or a continuous distribution would need to be used in place of the Poisson (for example, a Gamma distribution).

An alternative strategy, which may be preferred in some cases, is to absorb the TU-specific rescaling induced by normalization into the *λ* constant. In particular, let each TU have its own read-depth parameter *λ_j_* and set 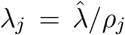, where 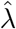 is the original (global) estimate of *λ*. In this way, the analysis can still make use of the original raw data **X**, rather than the normalized data **X**′, but the inferred rate parameter will effectively reflect the normalization transformation.

In tests of differences in the pause-release rate *β*, a second normalization may be needed. There are some indications that the degree to which RNAP accumulates in the promoter region may differ from one experiment to another, perhaps owing to differences in the concentration of factors that contribute to pause release. As a result, the data may display batch effects in pause peaks beyond those that can be explained by differences in the gene bodies. This problem could be addressed by a second round of normalization applied to the *X_k_* values after having normalized for differences in *s*′, as described above. As in the first case, this normalization could also be accommodated by altering the *λ* values per TU, but it would require the introduction of a second *λ_j_* that applied only to the pause peaks.

#### Replicates

In many applications, it is desirable to perform multiple replicates (biological or technical) of each individual experiment, to mitigate the noisiness of the sequence data and increase confidence in parameter estimates or hypothesis tests. Replicates can easily be accommodated in the model by working with a joint likelihood function that assumes independence across replicates but shares parameters that represent an underlying biological “truth” expected to be the same in each replicate. Typically, the key rate parameters—*α*, *β*, and *ζ*—will be shared, whereas technical parameters—such as *λ*—will be separate for each replicate.

With *R* replicates per time point, the likelihood function for the nonequilibrium case (cf. equation 4) generalizes to,

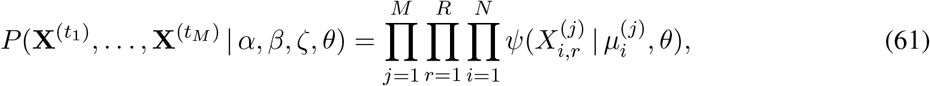

where 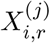 represents the read count for nucleotide *i* and time-point *j* as measured in replicate *r* and 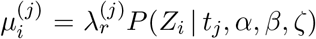 represents a shared “truth” across replicates for site *i* and time-point *j* that is separately normalized for each replicate *r* using read-depth parameter 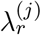.

In the case of the Poisson model at steady-state, the likelihood function reduces to (cf. equation 9),

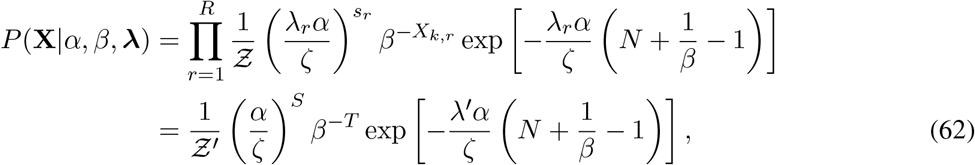

where 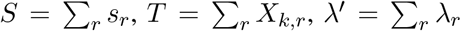, and where we absorb the terms that depend only on *λ_r_* in 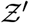, assuming that the *λ_r_* values will be pre-estimated. Thus, the likelihood function can be expressed in terms of sufficient statistics that are simply sums over replicates of the original sufficient statistics. Notice that this function would be identical if the read counts were simply pooled across replicates. Predictably, it leads to maximum-likelihood estimators of,

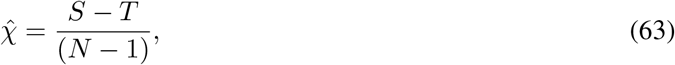

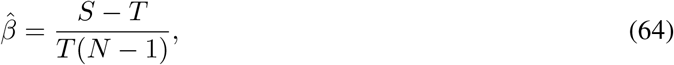

where, in this case, 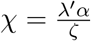.

The likelihood ratio tests and generalized linear model under the Poisson model can similarly be expressed in terms of these new aggregate sufficient statistics.

## Discussion

In this article, I have outlined a new probabilistic modeling framework for the analysis of nascent RNA sequencing data. This framework allows for a unified treatment of many problems of interest in the study of eukaryotic transcription, including estimation of the rates of initiation, elongation, and pause-release. It leads naturally to likelihood ratio tests for differences in rates, and it has extensions to Bayesian inference and a generalized linear model for evaluating covariates of elongation rate. It can also be adapted to capture the phenomenon of steric hindrance of initiation from paused polymerases at steady state. In addition, several issues of practical importance in the analysis of real data—such as accomodating differences in library size, batch effects, and experimental replicates—can be readily addressed in this framework.

A number of machine-learning and statistical methods already exist for analyzing nascent RNA sequencing data [6, 8, 12, 13], so it is reasonable to ask what is gained by addressing these problems using the generative probabilistic model described here. In my view, this generative model has several key advantages. First, the model does allow estimation of some quantities that have been out of reach of most previous methods, such as the relative rates of pause-release and initiation/elongation. These quantities have been indirectly characterized by more heuristic methods—as when pausing is quantified using the pause index—but it has been difficult to know precisely how to interpret these measures. Second, the model can be fitted to the raw read counts in a completely unsupervised manner, with no need for labeled training data. This property is important because, while high-quality, independent training data is available for some applications (e.g., [8]), no such data is available for others, including estimation of pause-release or elongation rates, or tests for differences in rates. Third, because the model allows several distinct problems to be addressed in a unified, coherent manner, it allows various parameter estimates and hypothesis tests to be interpreted jointly, without the concern that separate estimates might reflect different biases or different modeling assumptions—as when, say, applying one machine-learning method to estimate elongation rates and a second, completely separate method to characterize promoter-proximal pausing or correlates of elongation rate. Finally, by directly describing the actual process by which RNAP moves along the DNA template, the model potentially leads to new insights into this process, beyond simply providing predictions of quantities of interest, as a discriminative machine-learning method would. Altogether, the generative model both opens up new applications and provides a new perspective on the fundamental process of eukaryotic transcription.

The model does require a number of simplifying assumptions, some of which may not hold in all settings. For example, it assumes that RNAPs that initiate transcription gradually make their way all the way along the DNA template to the termination site, with negligible rates of premature termination. The frequency with which premature termination occurs is debated in the literature, with some studies suggesting it occurs at low rates across most genes [16] and others indicating higher rates [28, 29]. My hope is that this phenomenon occurs at low enough rates across most of the TU (with the possible exception of locations near the promoter) that it will have at most a minor impact on the rate estimators and hypothesis tests described here, but it could result in appreciable biases. The challenge in this setting is that, if premature termination were added to the model, its rate would be confounded with the initiation and elongation rates. I do not see a way of accommodating it in this framework except by conditioning on independent estimates of its rate.

As noted throughout the article, another major assumption is that each RNAP molecule moves independently along the DNA template, without interference from other molecules. This “no collisions” assumption should be reasonable in the regime where the density of RNAPs per TU per cell is low, as it appears to be on average. However, this assumption is likely violated frequently upstream of the pause peak, especially when the intitiation rate is high and/or the pause release rate is low. I have shown that the special case of collisions upstream of the pause site can be addressed at steady-state, within the framework of the Markov model, by taking advantage of the limited space for RNAP molecules in this region and assuming that the primary consequence of collisions is an effective decrease in the initiation rate. That is, I assume, to a first approximation, that either an RNAP is or is not present upstream of the pause site, and if it is, it will block initiation by new RNAPs. I have argued that, if indeed most collisions occur in this small local region, this adapted Markov model may be adequate for many purposes. But it is still unclear how frequently collisions occur downstream of the pause site, and more work will be needed to explore this case. In particular, high levels of temporal variation in the initiation rate (transcription “bursts”, e.g., [30]) could conceivably produce many more collisions than average rates would suggest. I have argued that the steady-state solution for the *ℓ*-TASEP recently introduced by Erdmann-Pham et al. [20] could be used in place of the stationary probabilities from the Markov model to accommodate collisions in a more general way. Still, as I understand it, this model considers pairwise interactions between RNAP molecules but ignores higher-order interactions, so Monte Carlo methods or other approximations would presumably be needed for a fully general treatment of the issue.

The assumption of a Poisson distribution for read counts leads to a great deal of mathematical simplicity, and I have therefore relied heavily on this version of the model. I see three major points in favor of the Poisson model. First, it leads to closed-form expressions that are easy to interpet, and help to develop intuition for how the data drive parameter estimates and hypothesis tests. Second, it provides a convenient and rapid means for performing many computations of interest, without the need for complicated, slow, and error-prone software for numerical optimization. Third, it is likely that, even when the Poisson assumption is unrealistic (as when read counts are clearly overdispersed), it will nevertheless have limited impact on many of the estimators or tests of interest, particularly if care is taken in post-processing and interpretation. For example, in LRTs, one Poisson model is compared with another, and if log likelihood ratios are interpreted with respect to an empirical null distribution, the effects of overdispersion should be considerably mitigated. On the other hand, if it does prove necessary to discard the Poisson model, the negative binomial model is still relatively convenient to work with, despite requiring numerical optimization. Notably, the fully general framework would potentially allow the use of distributions for read counts that allowed not only for overdispersion but also for autocorrelation along the genome sequence, such as a Gaussian process model or hidden Markov model. More work will be needed to determine whether the use of more complex and computationally intensive generating distributions is justified.

It is worth noting that the analysis of real nascent RNA sequencing data requires addressing a number of practical issues that have been glossed over in this article. For example, the model addresses uncertainty in the pause site in a reasonably general way, but it ignores uncertainty in the TSS, which can lead to a “smear” of read counts at the 5′ end of TUs and/or to a mixture arising from alternative TSSs at considerable distance from one another. We have recently introduced a program, called DENR [31], that attempts to deconvolve the contributions of multiple overlapping pre-RNA isoforms, and it could be used to preprocess the data. However, even with DENR, some filtering of the data is typically still necessary to ensure a clean signal. I have also largely avoided the issue of the termination site in this article, because we find that it is quite difficult to pinpoint precisely in real data, owing to transcriptional run-on and variability across cells. For many purposes it may be adequate to omit the signal near the 3′ end of the TU, as I have assumed, but in others it may be desirable to model this portion of the gene. In general, different inference problems will require different strategies for preprocessing and filtering.

Overall, I believe that the combination of a continuous-time Markov model and a generalized generating distribution for read counts, as outined in this article, provides a flexible and powerful modeling framework for nascent RNA sequencing data. The version of the model described here does make use of a number of simplifying assumptions, some of which may need to be relaxed over time. Nevertheless, the basic framework described here should, if nothing else, be useful as a starting point for further model and algorithm development.

## Acknowledgments

I thank Charles Danko, Yifei Huang, Yixin Zhao, Lingjie Liu, and Noah Dukler for helpful discussions, and Yixin Zhao for help with figure preparation.

